# Neuronal rhythms orchestrate cell assembles to distinguish perceptual categories

**DOI:** 10.1101/191247

**Authors:** Morteza Moazami Goudarzi, Jason Cromer, Jefferson Roy, Earl K. Miller

## Abstract

Categories are reflected in the spiking activity of neurons. However, how neurons form ensembles for categories is unclear. To address this, we simultaneously recorded spiking and local field potential (LFP) activity in the lateral prefrontal cortex (lPFC) of monkeys performing a delayed match to category task with two independent category sets (Animals: Cats vs Dogs; Cars: Sports Cars vs Sedans). We found stimulus and category information in alpha and beta band oscillations. Different category distinctions engaged different frequencies. There was greater spike field coherence (SFC) in alpha (∼8-14 Hz) for Cats and in beta (∼16-22 Hz) for Dogs. Cars showed similar differences, albeit less pronounced: greater alpha SFC for Sedans and greater beta SFC for Sports Cars. Thus, oscillatory rhythms can help coordinate neurons into different ensembles. Engagement of different frequencies may help differentiate the categories.

## Introduction

The prefrontal cortex (PFC) is a key player in cognitive control (Miller and Cohen, 2001; Blackman et al., 2016; Goodwin et al., 2012; Helfrich and Knight, 2016) and as such, needs access to top-down information that groups experience into meaningful abstractions such as categories and rules (Miller and Cohen, 2001; Seger and Miller, 2010), There are numerous examples of categories reflected in spiking activity in the PFC and elsewhere (Antzoulatos and Miller, 2011, 2016, Cromer et al., 2010, 2011;Ferrera et al., 2009; Freedman and Assad, 2006, 2016, Freedman et al., 2001, 2003; Hampson et al., 2004; Roy et al., 2010). Individual PFC neurons (as well as those in parietal cortex) do not encode single categories, however. They are multifunctional and participate in multiple categorical decisions. For example, in a previous study of ours (Cromer et al., 2010), we found that the majority of PFC participated in two different, unrelated, category distinctions (Cats vs Dogs or Sports Cars vs Sedans). Likewise, many LIP neurons encode both shape and motion categories (Fitzgerald et al., 2011).This multifunctionality seems to be key to PFC’s role in high-level cognition. It reflects “mixed-selectivity” (Rigotti et al., 2013). That is, neurons that are sensitive to many different non-linear combinations of tasks. The result is a high dimensional workspace in which function is distributed among many neurons and many neurons participate in many functions (Rigotti et al., 2013). This fluidity may provide the infrastructure for cognitive flexibility (Duncan and Miller, 2002, 2013) and adds the computational horsepower needed for complex behavior (Fusi et al., 2016; Rigotti et al., 2013). But it raises the question of how a unique categorical decision is read out of a workspace in which multiple category representations are intermixed on the neuron level.

One possibility is that information about context (i.e., which decision is currently relevant) allows the multidimensional representation to be reduced to a simple linear discrimination (Fusi et al., 2016; Rigotti et al., 2013). Another contribution may come from oscillatory dynamics. An ensemble unique to a single categorical decision can be carved out of a mixed selectivity shared workspace if neurons of the same ensemble synchronize their rhythms (Fries, 2005). Indeed, there are examples of rules and categories reflected in unique patterns of alpha/beta rhythms in the PFC and between the PFC and striatum and parietal cortex (Antzoulatos and Miller, 2014, 2016; Buschman et al., 2012; Stanley et al., 2016).

Here, we tested whether oscillatory rhythms can reflect unique categorical decisions in the PFC. We re-examined data from Cromer et al (Cromer et al., 2010). Monkeys performed a delayed-match-to-category (DMC) task with two independent stimulus sets (Animals vs Cars). On each trial, monkeys had to distinguish between categories within each of these stimulus sets (Animals: Cats vs Dogs and Cars: Sport Cars vs Sedans). We examined the strength (power), frequency, and coherence of oscillations of local field potentials (LFPs) and their interactions with spiking activity (Spike-Field Coherence, or SFC). We found that each independent stimulus set (Animals vs Cars) produced increases in oscillatory power at different times and in different frequency bands (alpha vs beta). Within each stimulus set, different categories (Cats vs Dogs or Sport Cars vs Sedans) were reflected in increases in spike field coherence (SFC) at different ranges of frequencies.

## Results

### Behavior

Two monkeys performed well (over 95% correct) in the DMC task (Figure 1A–C, for more detail see Cromer et al., 2010) for both category sets. We used correct trials for neurophysiological analyses. Reaction time (RT) was significantly longer for Car category judgments than Animal category judgments (Figure 1D, Wilcoxon rank-sum test with α=0.01 rejected the null hypothesis). Within each category set, the median reaction time was slightly longer for Sedans than Sports Cars and for Dogs than Cats (Figure 1E, Wilcoxon rank-sum test with α=0.01 rejected the null hypothesis). This suggests that Cars were more difficult to categorize than Animals and, more specifically, task difficulty increased depending on whether the sample stimulus was a Cat (lowest difficulty) to Dog to Sports Car to Sedan (highest difficulty) trials (Figure 1E).

**Figure 1.**
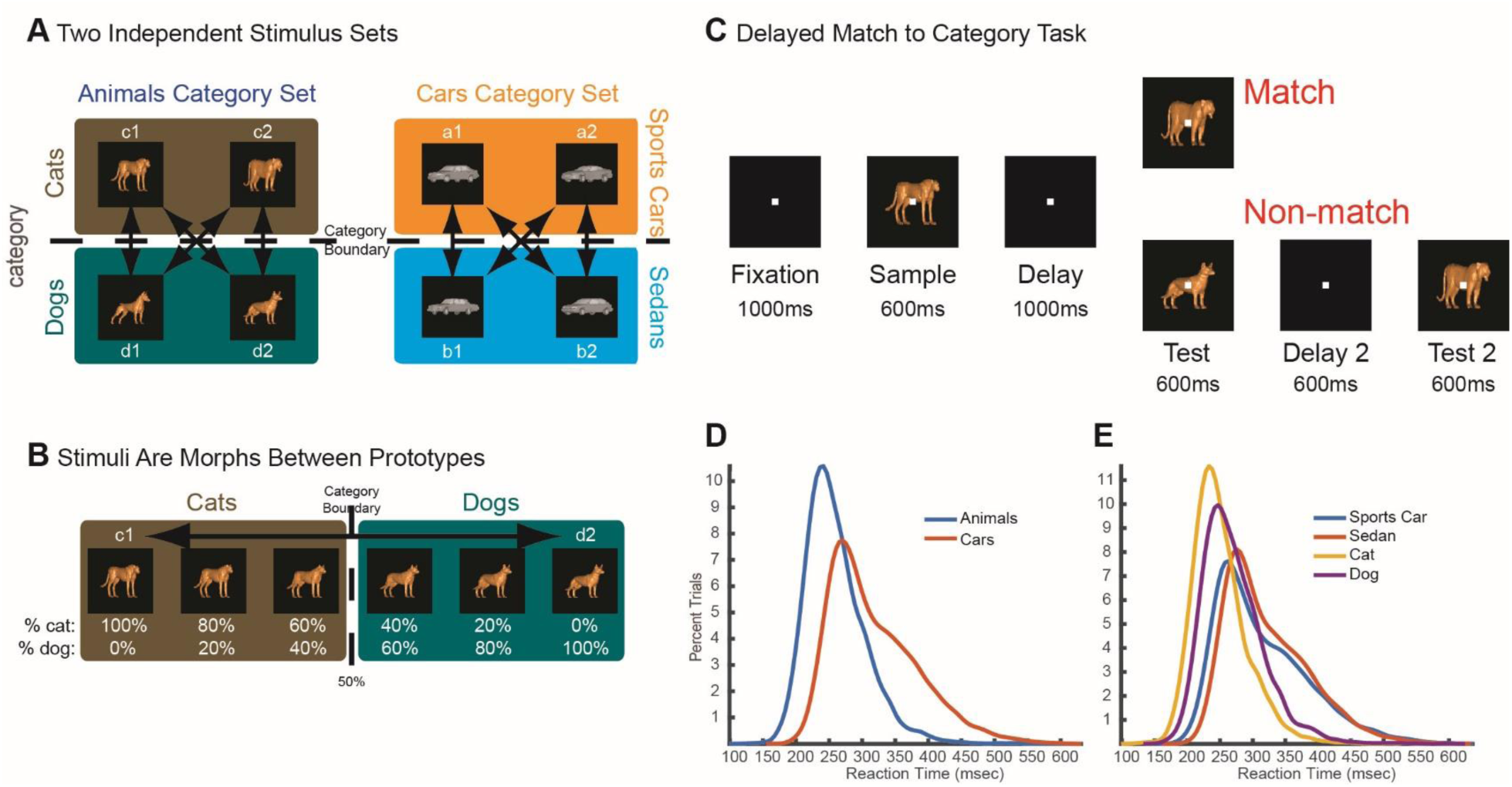
Task Design and Behavioral Performance. (A) Stimuli drawn randomly from animal or car category sets. Each set had two different categories (the animal set had Cats/Dogs and the car set had Sports Cars/Sedans). Both sets contained four prototype images (two from each category as shown). (B) Sample images were generated by morphing between prototypes along four morph lines. For example, if a sample belonged to the animal category set, sample images were morphed between Cat prototype c1 and Dog prototype d2. If sample images were comprised of greater than 50% of one category prototype, they were labeled as a member of that category. (C) Schematic of delay match to category task (DMC). In each task trial, after a 600 ms fixation period, one of the two possible morphs (either a Cats/Dogs morph or a Sports Cars/Sedans morph) was shown (sample image) followed by a 1 second delay and a test stimulus. If the sample image and test stimulus belonged to the same category (match trials), the monkeys released a lever. Alternatively, if they were non-matched, the monkeys continued to hold the lever for another 600 ms delay which was followed by a match test stimulus. (D) Blue and red curves indicate the empirical probability density function (ePDF) of RTs for animal and car category sets, respectively. (E) ePDF of RTs for each category (Cats, Dogs, Sports Cars, and Sedans).

### Average population LFP power and coherence distinguished stimulus sets, but not categories

We found that local field potential (LFP) power in the PFC was different for the Animals vs Cars stimulus sets. We defined an estimator as the difference of time-frequency power between the two category sets. We used the z-statistic, calculated for each electrode, to determine the statistical significance of our estimator. The difference in the time-frequency representation of the LFP signal was estimated for Animals vs Cars. A null distribution for this difference was estimated by randomly shuffling the trial labels (Animals, Cars) between trials. Finally, the original mean difference was subtracted from the mean of the shuffled differences and normalized by the standard deviation of the shuffled population.

This revealed stimulus set (Animals vs Cars) selectivity in LFP power that differed in frequency and time. During the sample epoch (150–600 ms after sample onset) there was greater power for Cars in the 14-21 Hz (beta) frequency band for ∼32 percent of the electrodes. This can be seen in Figure 2A which plots the average z-score of induced power differences between Animals and Cars across all electrodes as a function of time and frequency. Blue indicates greater average power for Cars, red for Animals. Of the electrodes that showed significant differences in power to Cars vs Animals during the sample epoch, nearly all showed greater power for Cars (Figure 2B, binomial test, p<10^−15^).

**Figure 2.**
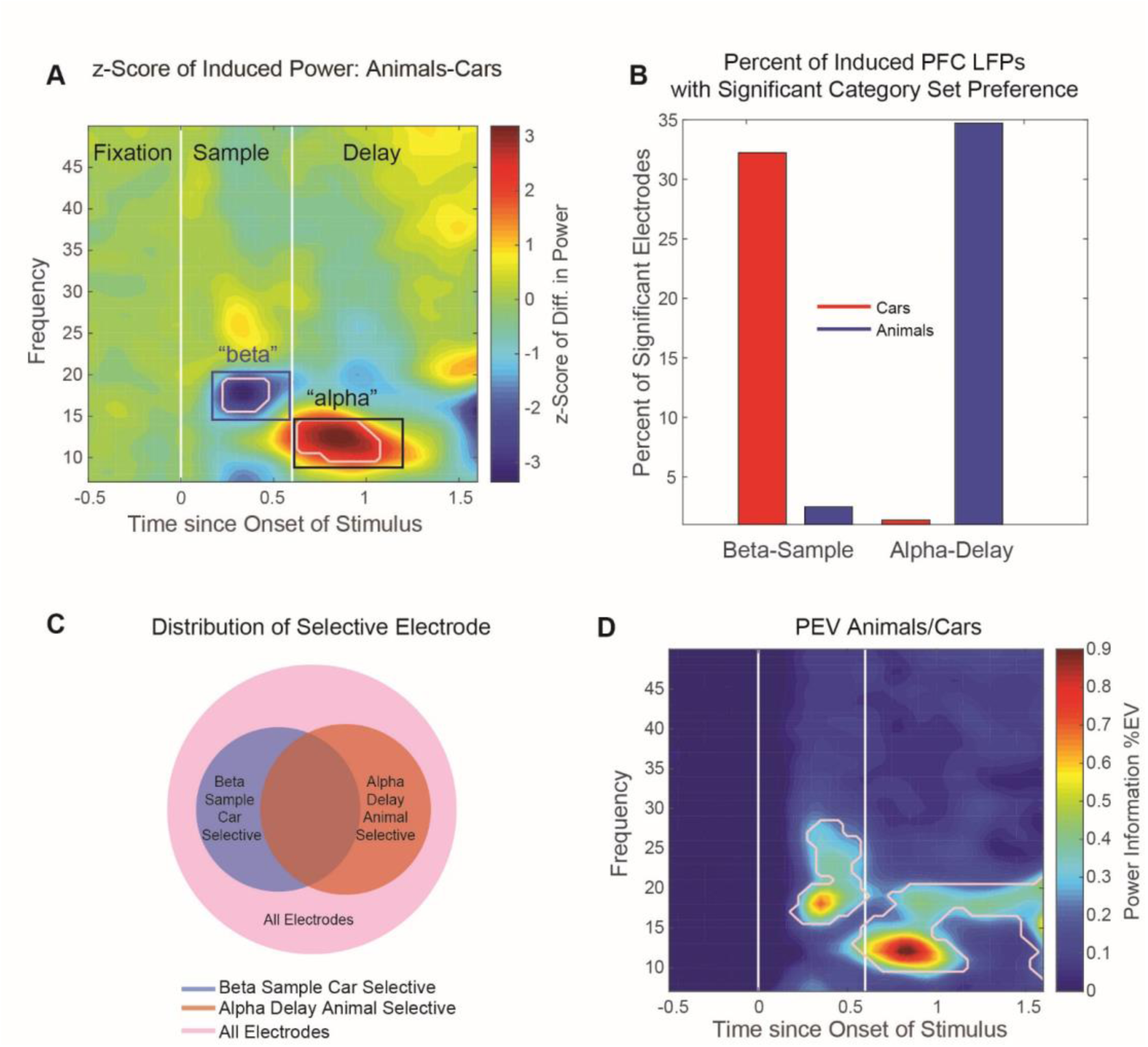
Independent Stimulus Set Selective LFP Power in PFC. (A)LFP power in PFC shows independent stimulus set selectivity. The color-coded image shows z- statistic of differences in LFP power between animal and car category sets (horizontal axis stands for time in ms from onset of sample image and vertical axis stands for frequency in Hz). In two time- frequency regions (shown with white contours) during sample and delay periods, LFP power was selective (pFDR<0.01) for car and animal category sets, respectively (p values were corrected for multiple comparisons using the false discovery rate, pFDR). (B) Percentage of LFP electrodes with a significant category selectivity in the two time-frequency regions, 150-600 ms and 14-21 Hz (blue outline) and 600-1300 ms and 9-14Hz (black outline). (C) Venn diagram of all LFP electrodes: all electrodes (pink), animal selective during delay (red), and car selective during sample (blue). (D) LFP power information about animal vs car category sets. The maximal information about independent category sets were located in the same time-frequency regions (white contours show TF regions with significant information about Animals/Cars, Wilcoxon signed-rank test, Bonferroni corrected p<10^−12^).

By contrast, in the delay epoch (600-1300 ms after sample onset), there was greater power for Animals in the 9-14 Hz (alpha) frequency band for ∼35 percent of the electrodes (red colors, Figure 2A). Most of the electrodes that showed significant differences in power between Cars and Animals during the delay, showed greater power for Animals (Figure 2B, binomial test, p<10^−15^). A subset of the electrodes (∼17%) showed both effects: Greater beta power for Cars during the sample and greater alpha power for Animals during the delay (Fig 2C).

We used an unbiased percent explained variance measure (ωPEV, see Methods section for more detail) to estimate the information about the Cars vs Animals in LFP power (Figure 2D). The time-frequency profile of information about Cars vs Animals was similar to the LFP power selectivity described above (Figure 2A). However, while average LFP power distinguished between Cars and Animals stimulus sets, it did not distinguish between the categories within each set (i.e. Cat vs Dog or Sports Car vs Sedan) (Supplementary materials Figure 1S A-D).

We conducted a similar analysis using synchrony between electrodes. The results almost mirrored that of the differences in power, but mainly in the beta band (Figure 2S). Like average power, population average LFP synchrony distinguished between Animals and Cars stimulus sets, but did not distinguish between categories within each stimulus set (Supplementary materials Figures 3S A-D).

### Coherence between individual electrode pairs was category selective

Above, we showed that *population average* power and coherence distinguished between stimulus sets but not the categories within them. However, we did find category selectivity within stimulus sets when we examined coherence between *individual pairs* of electrodes.

For LFPs from each possible pair of simultaneously recorded electrodes, we determine which showed significantly greater coherence for one category vs its alternative (Cats vs Dogs, Sports Cars vs Sedans selective, see Materials and Methods). We found category selectivity in electrode pair coherence in a similar frequency range to the average power effects (above). Figure 3A, B shows this selectivity expressed as the percentage of electrode pairs that showed significant differences in coherence for Cats vs Dogs (Fig 3A) and Sports Cars vs Sedans (Fig 3B). The selectivity was maximal at ∼9-25 Hz during both sample and delay epochs but overall stronger during sample presentation (Figure 3A, B). Figure 3C, D shows the average of absolute z-score of proportion of electrode pairs that showed significant differences in coherence.

**Figure 3.**
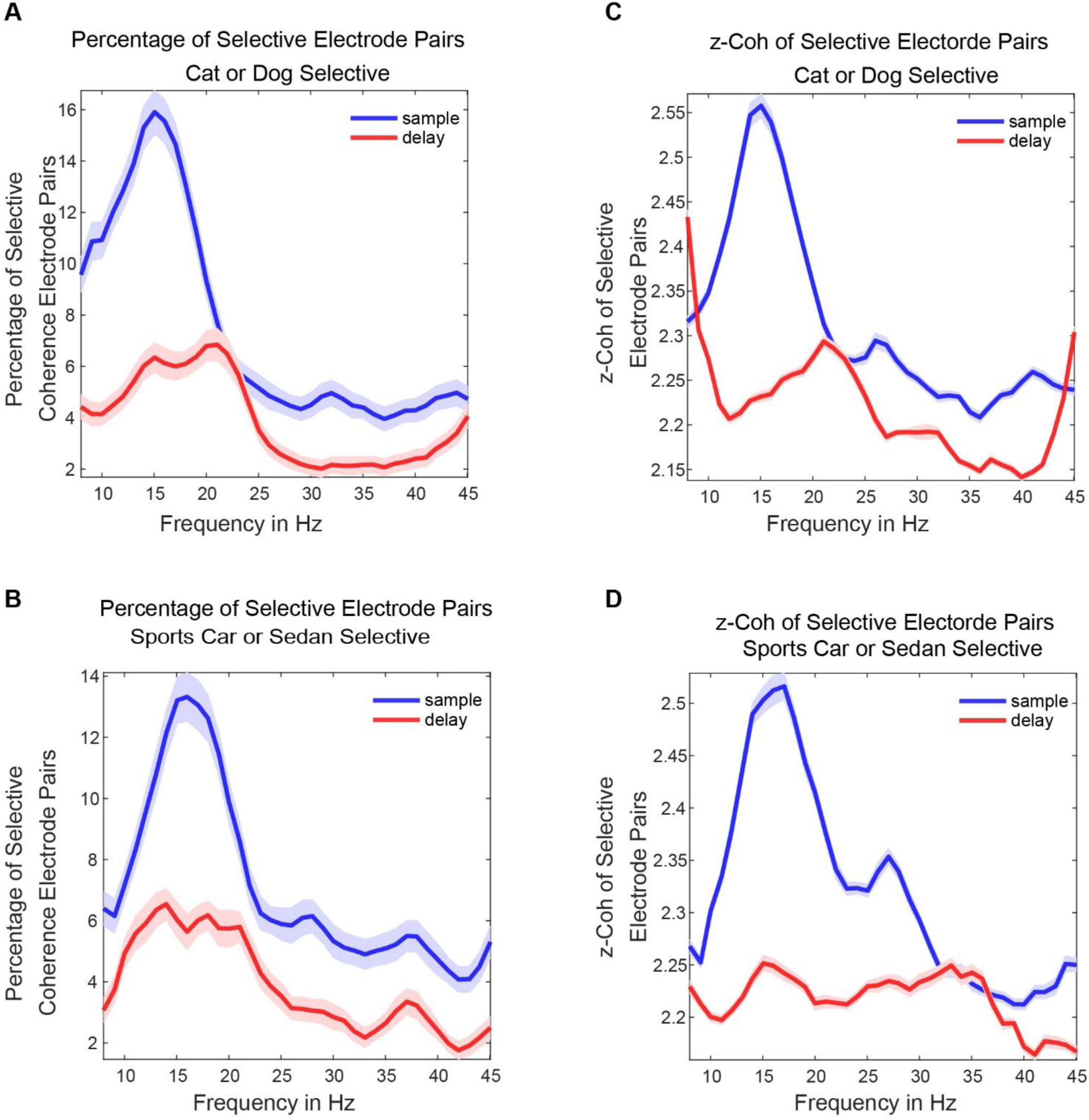
Selectivity Analysis for Coherence Pairs within Category Sets. (A), (B) Curves show percentage of significant electrode pairs with category distinction selectivity (Cats vs Dogs and Sports Cars vs Sedans) during sample (blue) and delay (red) epochs. (C), (D) Curves show the absolute value of the z-score of selective electrode pairs with category distinction selectivity (Cats vs Dogs and Sports Cars vs Sedans) during sample (blue) and delay (red) epochs.

### Spike-field coherence distinguished between independent stimulus sets and the categories within them

It has been suggested that oscillatory rhythms may help organize the spiking activity of ensembles of neurons (Buschman et al., 2012; Lundqvist et al., 2016). Thus, we next examined spike-field coherence between LFPs and single isolated neurons. To maximize our dataset, we averaged across the sample and delay epochs. Once again, we found effects in the alpha-beta range. Figure 4 shows spike-field coherence averaged across all electrodes and spikes. The population average SFC reached a higher peak for the Cars than Animals stimulus set in the beta range ∼16-22 Hz (Fig 4A, dotted line shows significant differences with pFDR<0.01). But note that in the alpha range (∼8-12 Hz), there was greater average SFC for Animals than Cars.

**Figure 4.**
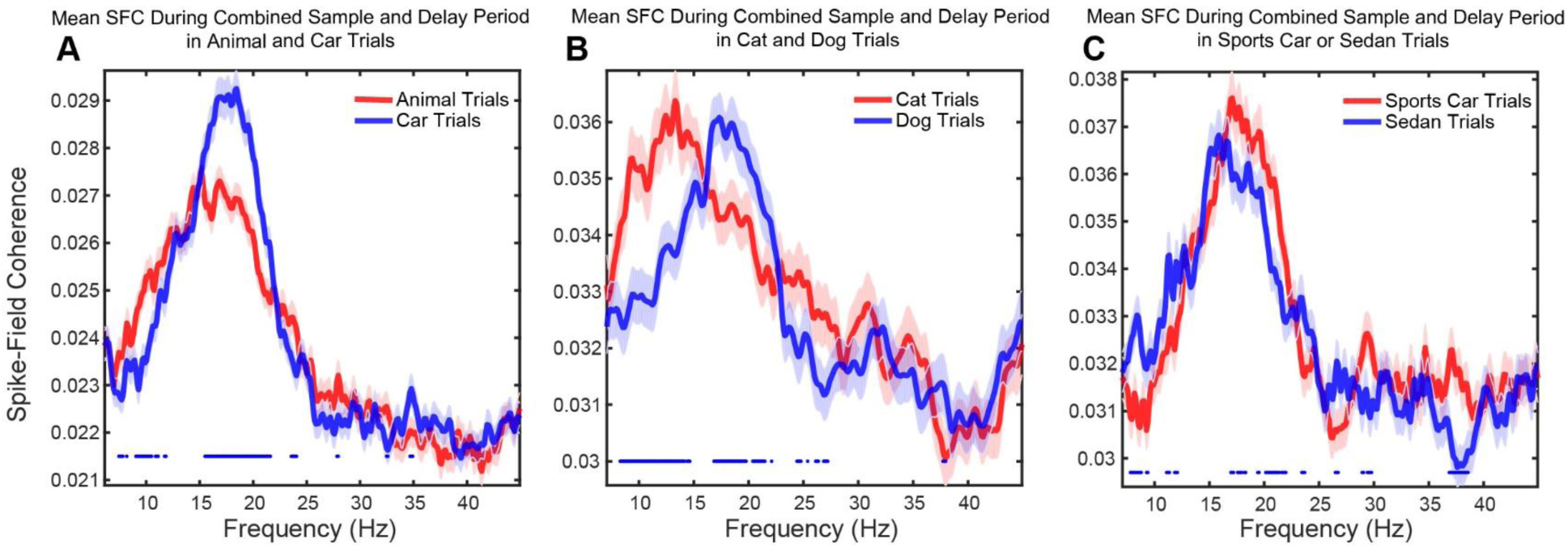
SFC Selectivity for Independent Stimulus Sets and Categories. SFC as a function of frequency during the combined sample and delay epoch. Blue and red curves show average SFC between all LFPs with all units in the following independent stimulus sets and categories. The dotted lines show statistical differences between the blue and red curves, t-test pFDR<0.01. (A) Car and Animal trials, respectively. (B) Dog and Cat trials, respectively. (C) Sedan and Sports Car trials, respectively.

This difference in population average SFC for Animals and Cars seems to be due to frequency differences in SFC for different categories within the stimulus sets. Different categories showed greater SFC at different frequencies. This was especially true for the Animals (Fig 4B). Note that the average population SFC was higher for Cats than Dogs at ∼ 8-14 Hz but higher for Dogs than Cats at ∼16-22 Hz (Fig 4B). There was also significantly higher SFC for Cats than Dogs around 25 Hz (Fig 4B), but those differences were not as pronounced as the others. For the Cars, similar differences were observed, albeit less marked. The average population SFC was significantly higher for Sedans than Sports Cars at ∼ 8-12 Hz and significantly higher for Sports Cars than Sedans in ∼16-23 Hz (Figure 4C,). We also asked whether SFC between *individual* pairs of electrodes (rather than the population average) can distinguish categories. We estimated the percentage of SFC pairs selective for categories as a function of frequency (for more details see Materials and Methods). This revealed category selective SFC in the ∼8-25 Hz frequency range (Figure 4S A-D).

## Discussion

We found that category and stimulus information was reflected in the oscillatory dynamics between local field potentials (LFPs) and between LFPs and spikes in the lateral PFC. LFPs showed greater average power in the beta (∼14-21 Hz) band for the Car stimulus set and greater average power in alpha (∼9-14 Hz) band for the Animal stimulus set. But neither population average power nor average synchrony were able to distinguish between the categories within each stimulus set (Cats vs Dogs or Sports Cars vs Sedans). Instead, category selectivity was seen on a more specific level:Changes in coherence between individual pairs of electrodes and in differences in spike-field coherence (SFC) between spikes and LFPs. There was significantly greater average SFC for Cats in the alpha range and for Dogs in the beta range. Similarly, there was greater average SFC for Sedans in alpha and for Sport Cars in beta. These differences were more pronounced for Animals than Cars, which may be due to the monkeys having more experience categorizing Cats vs Dogs than Sport Cars vs Sedans. Indeed, behavioral reaction time was shorter for Animals than Cars suggesting the former was an easier distinction. Thus, neural synchrony might act as a “glue” for coordinating cell assemblies at distinct frequencies. It could use the frequency dimension as an extra workspace to dissociate the categories.

Oscillatory synchrony has been shown to be involved in the formation of functional networks in cognition as well as different functional brain disorders (Buschman et al., 2012; Fries, 2005; Llinás et al., 1999; Moazami-Goudarzi et al., 2008; Spitzer and Haegens, 2017; Uhlhaas and Singer, 2006; Voytek et al., 2015; Womelsdorf and Everling, 2015; Womelsdorf and Fries, 2007). All three measures that we used, power, coherence, and SFC, capture different phenomena to some extent but they are all in consensus that the alpha and beta frequency bands are playing a role in representing category information. This is consistent with recent studies showing that categories and rules are reflected in alpha/beta coherence between different recording sites within the PFC (Antzoulatos and Miller, 2016; Buschman et al., 2012; Stanley et al., 2016), between PFC and striatum (Antzoulatos and Miller, 2014), and between PFC and posterior parietal cortex (Antzoulatos and Miller, 2016). In fact, we found that PFC beta synchrony was better at filtering out irrelevant category information than spiking (Antzoulatos and Miller, 2016), suggesting that beta synchrony was a more “pure” top-down signal than spike rates. By contrast, higher, gamma frequencies (>40 Hz) have been associated with bottom-up sensory inputs (Bastos et al., 2015; Buschman and Miller, 2007; van Kerkoerle et al., 2014; Lundqvist et al., 2016). Thus, our results fit with the hypothesis that alpha/beta rhythms provide an infrastructure for top-down cortical processing (Bastos et al., 2015; Buschman and Miller, 2007; Engel and Fries, 2010).

SFC, in particular, has been related to functional recruitment of cell assemblies (Wong et al., 2016) and beta oscillations have been associated with the dynamic formation of functional cell assemblies (Kopell et al., 2011). Thus, this range of frequencies may be particularly important for top-down information because it is well-suited for creating unique ensembles from the populations of mixed selectivity neurons that are critical for high-level cognition.

Indeed, category information has been found in brain areas where there is also evidence of mixed selectivity in spiking, such as the prefrontal cortex (Cromer et al., 2010; Freedman et al., 2001; Miller et al., 2002; Roy et al., 2010), parietal cortex (Fitzgerald et al., 2011), and hippocampus (Hampson et al., 2004). Our previous analysis of spiking activity from this data set (Cromer et al., 2010) showed that most PFC neurons that convey category information were “category generalists” in that they participated in two, unrelated category distinctions, a property consistent with mixed selectivity. Likewise, Freedman and colleagues showed that many of the same LIP neurons participate in shape categorization and direction of motion categorization. Thus, our results suggest that oscillatory dynamics can be used to form unique ensembles of categories in a higher dimensional mixed selectivity neural space where different task factors intermingle on the neuron level. It contributes to a body of evidence that neural ensembles -rather than single neurons- are the functional units of the brain (Eichenbaum, 2017; Yuste, 2015).

## Materials and methods

### Animals

Data were recorded from two adult rhesus macaques (one female, one male, *Macaca mulatta*) in accordance with the National Institutes of Health guidelines and the policies of the Massachusetts Institute of Technology Committee on Animal Care.

### Behavioral Task

Two monkeys were trained to perform a delayed match to category task for two independent category sets (Animals: Cats vs Dogs and Cars: Sports Cars vs Sedans). On a given trial, one stimulus from either stimulus set was shown as a sample, followed by a 1-second delay and a test stimulus. If the sample and test stimuli belonged to the same category (both Cats or Dogs or both Sports Cars or Sedans), the monkey was required to release a lever less than 600 ms after the test stimuli presentation (match trials) which resulted in reward and ended the trial, otherwise, (for nonmatch trials), the monkey had to continue holding the bar for another 1200 ms: 600 ms until the test stimuli turned off, followed by another 600ms delay period. Then, upon presentation of a second test stimulus, which always belonged to the same category as the sample image, the monkey was required to release the bar in less than 600 ms. Therefore, on every trial, in order to get a reward the monkey was required to make a correct response (see Figure.1 and Cromer et al., 2010 for more details).

## Recordings

We simultaneously recorded spiking and local field potential (LFP) activity in the lateral prefrontal cortex (lPFC) of monkeys performing a delayed match to category (for more details of recording see Cromer et al., 2010). Over 34 recording sessions, 517 isolated neurons and 363 electrodes were recorded in lPFC from both monkeys.

## Data analysis

Data were analyzed using custom MATLAB scripts (The MathWorks, Natick, MA) and the Chronux and Fieldtrip toolboxes (Mitra and Pesaran, 1999; Oostenveld et al., 2011). All analyses were performed on correct trials. In all analyses evoked activity was removed from the LFP. Thus, power, coherence spectra, and SFC were estimated on the induced LFP. Time-frequency representation of LFP power was estimated using a fast Fourier transform (FFT) applied on LFP data tapered by a 0.5 s Hanning window and in 0.05 s sliding steps in the time interval between −1 to 2 s from sample onset and in the 1-50 Hz frequency range. Synchrony was estimated by coherence (Phase locking value showed the same results as coherence, data not shown) between all simultaneously recorded possible pairs of electrodes. Significant differences in power and coherence between two independent category sets (Animals and Cars) or within each category set (Cats vs Dogs or Sports Cars vs Sedans) were inferred with a z-statistic, under the null hypothesis that there are no differences in power or synchrony between Animals and Cars or within category sets. The null distribution was generated by randomly shuffling the trial labels between two conditions at least 200 times. The mean and variance of this distribution was used to estimate the z-score of the main differences in power or coherence. An absolute value of the z-score greater than 1.96 corresponded to significant differences between the two conditions.

SFC was estimated using multitaper methods during the combined sample and delay epoch (1600 ms) with time-bandwidth product TW= 4 and K=7 tapers. Number of trials were equated for both conditions (to correct the biases due to the different number of trials per condition). If one condition had more trials, a random subset of trials equal to the minimum number of trials from other conditions was chosen. SFC was estimated for all simultaneously recorded Spikes and LFP that had not been recorded from the same electrode (distance of > 1mm) to exclude any coherence between LFP and Spike train that might be due to spike leakage to the LFP. To test population differences between SFC for different conditions we used t-tests and Wilcoxon sign-rank tests, which showed almost identical results. We applied the false discovery rate (FDR) (Benjamini and Hochberg, 1995) procedure to correct for multiple comparisons. To test SFC pairs selectivity, we used z-statistics. To this end, we permuted the label between conditions and estimated the SFC differences between the two conditions for each permutation (200 permutations). The mean and standard deviation of this null distribution was used to estimate the z-score of the main differences in SFC between the two conditions. Then SFC pairs were considered selective if their absolute z-score was greater than 1.96. A similar procedure was applied to estimate selectivity for coherence. Information was estimated using ω^2^ –statistics (Olejnik and Algina, 2003), which is a less biased statistic. To estimate information in coherence about category, coherence was estimated using the multitaper method with time-bandwidth product TW= 2 and K=3 tapers; for power the same parameters were used that we used to estimate time-frequency representation of LFP power. The significance of ω^2^ information about the categories in power and coherence was determined by a Wilcoxon signed-rank test and corrected for multiple comparisons using Bonferroni correction.

## Acknowledgements

We thank A. Bastos, A. Wutz, A. Khani, and M. Wicherski for comments on the manuscript, and E. Antzoulatos, I. Aganj, and S. Brincat for helpful discussions. This work was supported by NIMH R01MH065252 and the MIT Picower Institute Innovation Fund.

---

**Figure 1S.**
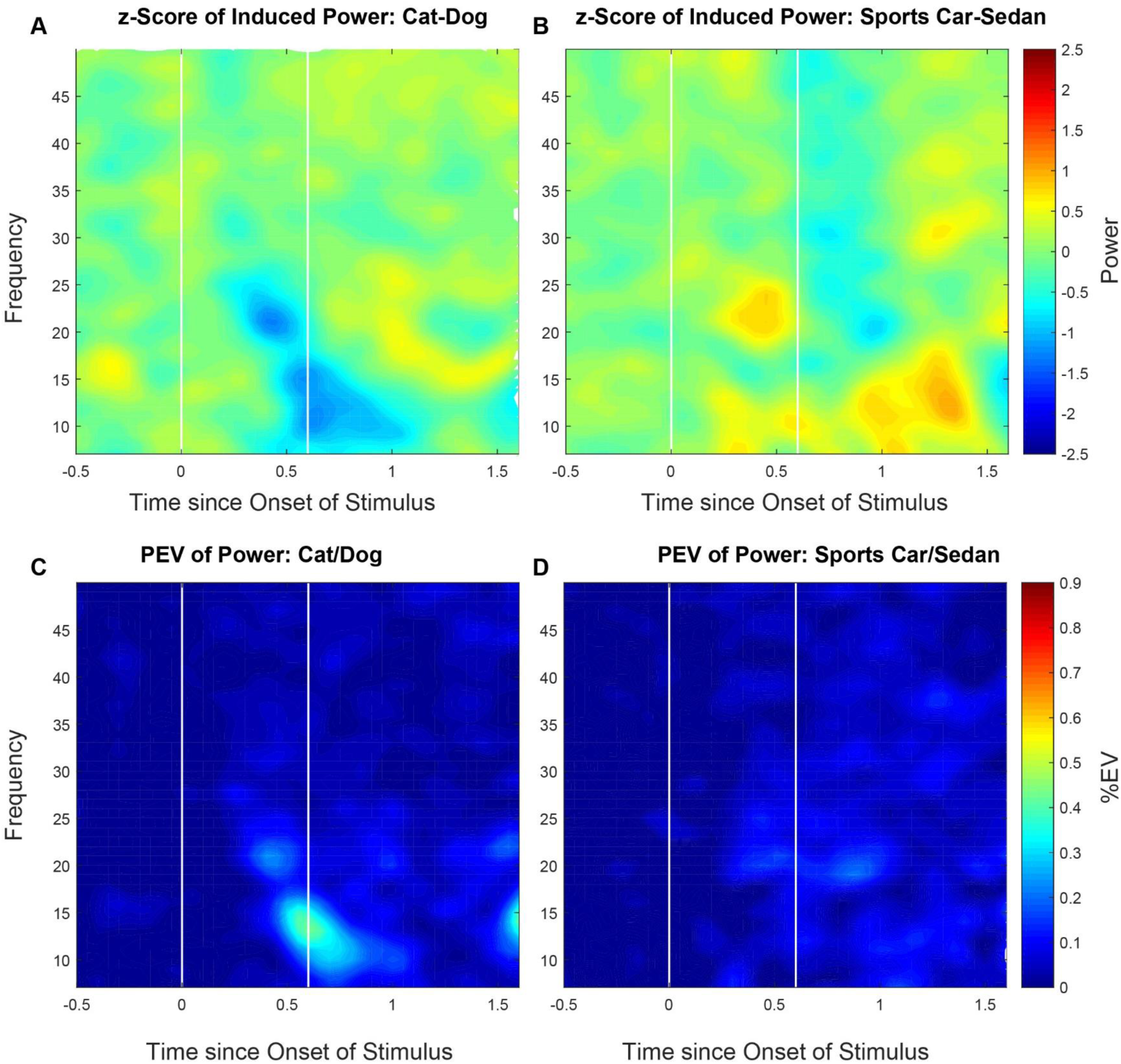
Average LFP Power Differences are Blind to Categories. (A), (B) Average z-statistic of induced LFP power difference between categories (Cats and Dogs, Sports Cars and Sedans). (C), (D) Information in power for category distinction (Cats vs Dogs, Sports Car vs Sedan).

**Figure 2S.**
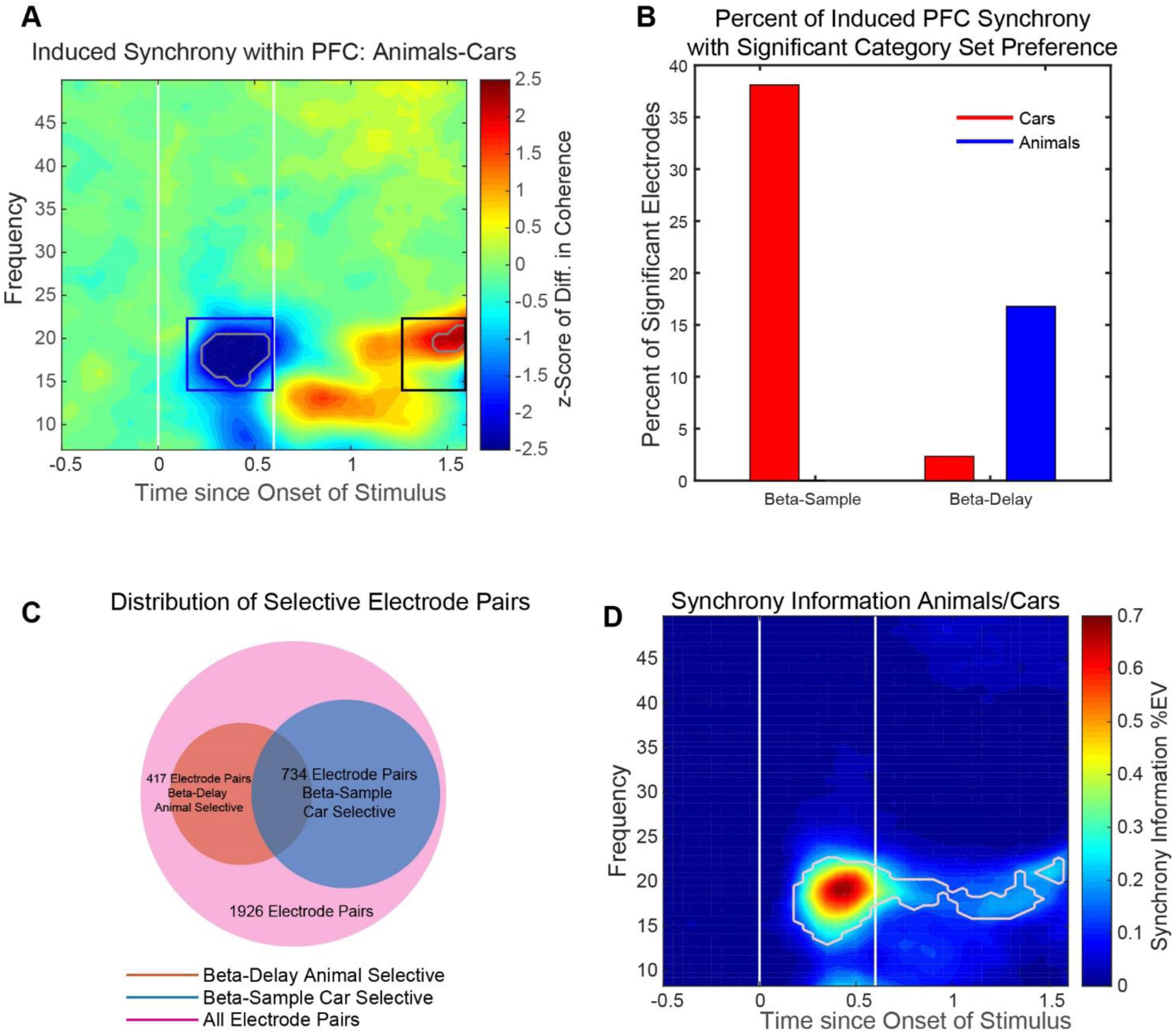
Independent Stimulus Set Selective LFP Synchrony in PFC. (A) Time-frequency LFP coherence in PFC shows independent stimulus set selectivity. Color-coded image shows z-statistic of differences in coherogram between animal and car category sets (horizontal axis stands for time in ms from onset of sample image and vertical axis stands for frequency in Hz). In two time-frequency regions (shown in white contours), during sample and delay periods, coherence was selective for car and animal category sets, respectively. (B) Percentage of LFP electrode pairs with significant category selectivity estimated in the two time-frequency regions, at 14-22 Hz and at 150-600 ms (blue outline) and 1300-1600 ms (black outline). (C) Venn diagram of all LFP electrode pairs: all electrodes (pink), animal selective during delay (red), and car selective during sample (blue). (D) LFP coherence information about animal vs car category sets. The maximal information about independent category sets was located in the same frequency band (14-22 Hz) during the sample and delay epochs (white contours show coherogram regions with significant information about Animals/Cars, Wilcoxon signed-rank test, Bonferroni corrected P<10^−15^).

**Figure 3S.**
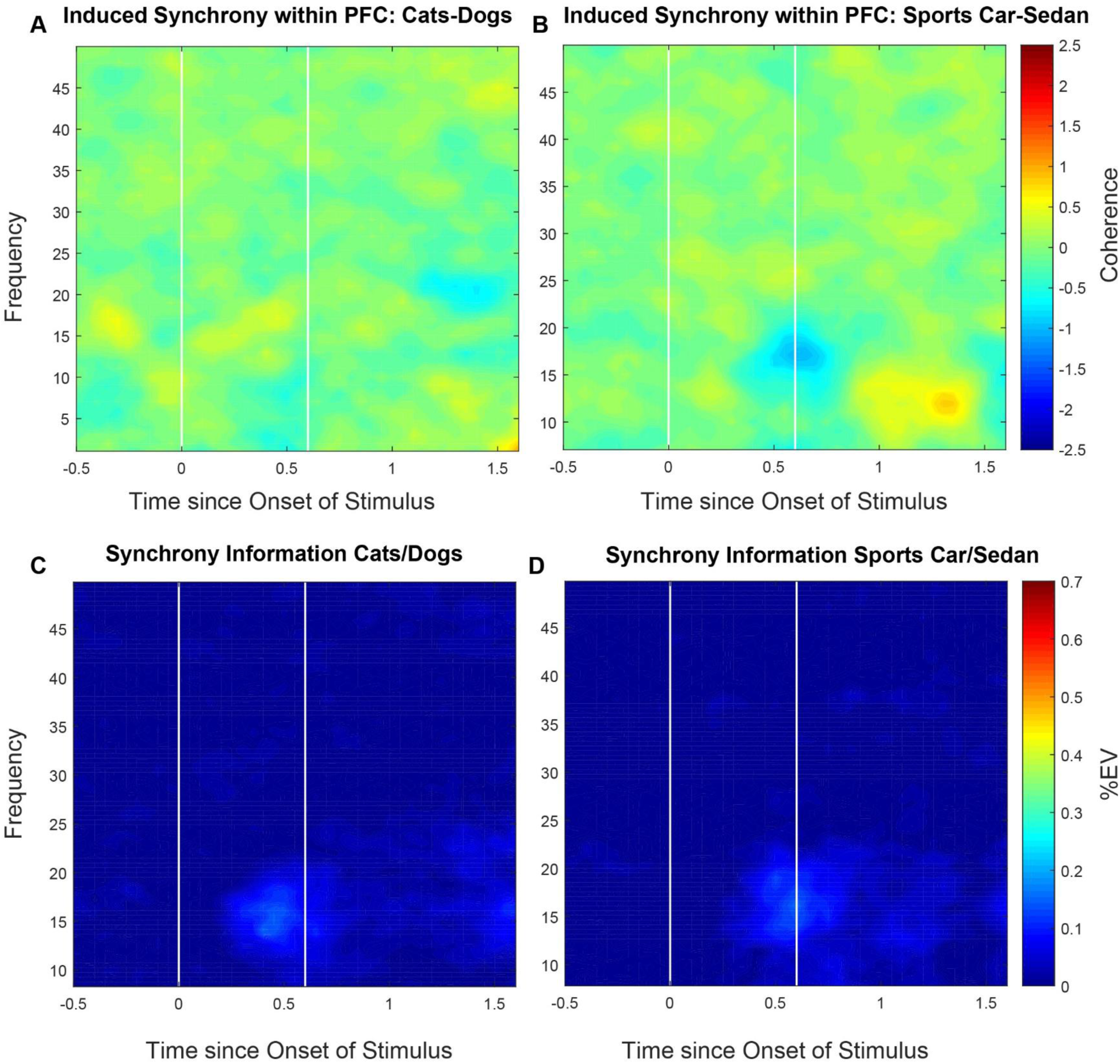
Average LFP Coherence is Blind to Categories. (A), (B) Average z-statistic of all PFC LFP pairs of induced coherence difference for both sets of categories (Cats and Dogs, Sports Cars and Sedans). (C), (D) Information in coherence for category distinction (Cats vs Dogs and Sports Cars vs Sedans).

**Figure 4S.**
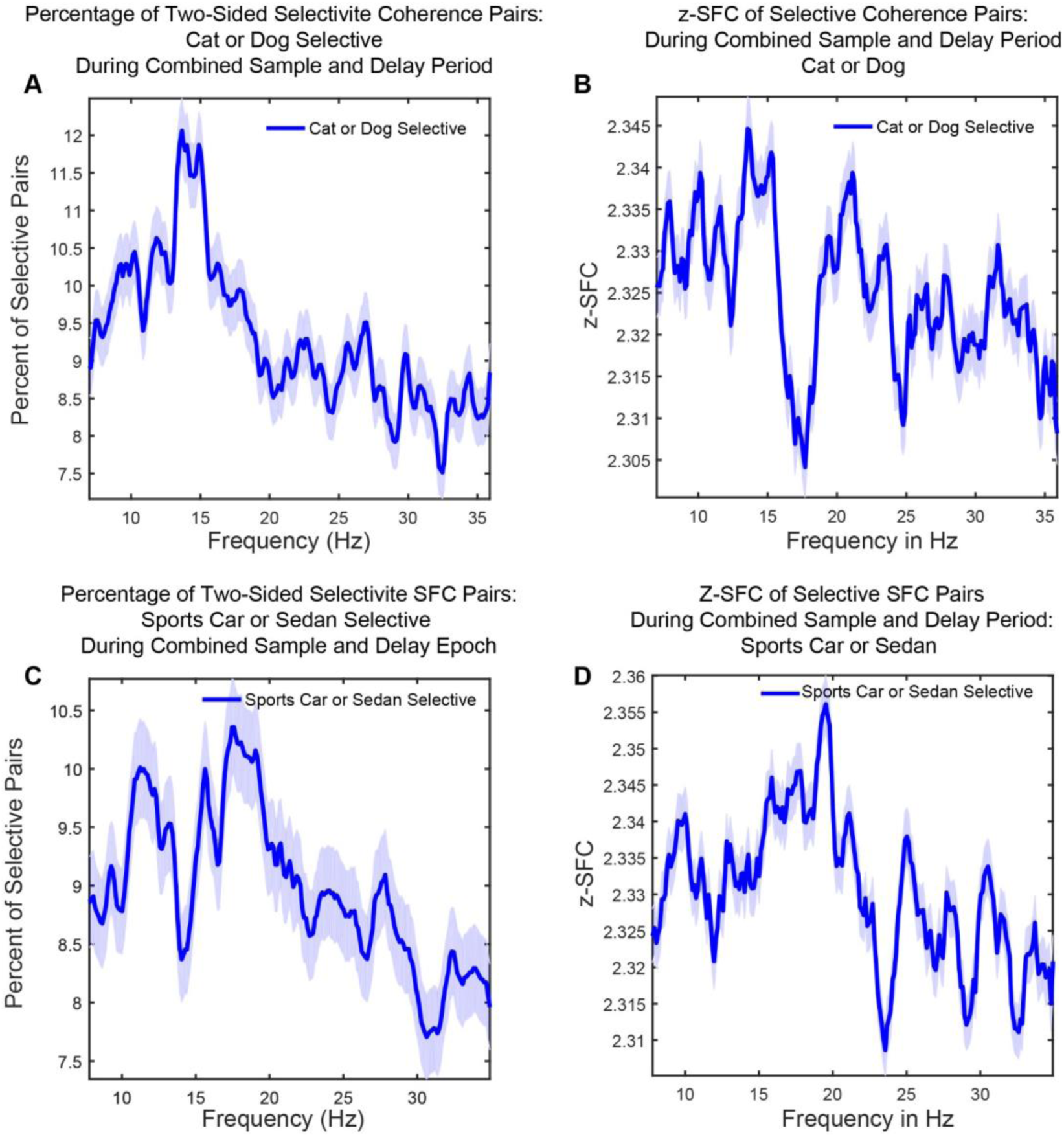
Selectivity Analysis of SFC Pairs for Categories. (A), (C) Percentage of significant SFC pairs with category distinction selectivity, Cats vs Dogs and Sports Cars vs Sedans, respectively. (B), (D) Absolute values of the z-score of significant SFC pairs with distinction selectivity within categories, Cats vs Dogs and Sports Cars vs Sedans, respectively.

## References

Antzoulatos, E.G., and Miller, E.K. (2011). Differences between neural activity in prefrontal cortex and striatum during learning of novel abstract categories. Neuron 71, 243–249.

Antzoulatos, E.G., and Miller, E.K. (2014). Increases in functional connectivity between prefrontal cortex and striatum during category learning. Neuron 83, 216–225.

Antzoulatos, E.G., and Miller, E.K. (2016). Synchronous beta rhythms of frontoparietal networks support only behaviorally relevant representations. Elife 5, e17822.

Bastos, A.M., Vezoli, J., Bosman, C.A., Schoffelen, J.-M., Oostenveld, R., Dowdall, J.R., De Weerd, P., Kennedy, H., and Fries, P. (2015). Visual areas exert feedforward and feedback influences through distinct frequency channels. Neuron 85, 390–401.

Benjamini, Y., and Hochberg, Y. (1995). Controlling the false discovery rate: a practical and powerful approach to multiple testing. J. R. Stat. Soc. Ser. B Methodol. 289–300.

Blackman, R.K., Crowe, D.A., DeNicola, A.L., Sakellaridi, S., MacDonald, A.W., and Chafee, M.V. (2016). Monkey Prefrontal Neurons Reflect Logical Operations for Cognitive Control in a Variant of the AX Continuous Performance Task (AX-CPT). J. Neurosci. 36, 4067.

Buschman, T.J., and Miller, E.K. (2007). Top-down versus bottom-up control of attention in the prefrontal and posterior parietal cortices. Science 315, 1860–1862.

Buschman, T.J., Denovellis, E.L., Diogo, C., Bullock, D., and Miller, E.K. (2012). Synchronous oscillatory neural ensembles for rules in the prefrontal cortex. Neuron 76, 838–846.

Cromer, J.A., Roy, J.E., and Miller, E.K. (2010). Representation of Multiple, Independent Categories in the Primate Prefrontal Cortex. Neuron 66, 796–807.

Cromer, J.A., Roy, J.E., Buschman, T.J., and Miller, E.K. (2011). Comparison of Primate Prefrontal and Premotor Cortex Neuronal Activity Visual Categorization.

Duncan, J., and Miller, E.K. (2002). Cognitive focus through adaptive neural coding in the primate prefrontal cortex. In Principles of Frontal Lobe Function, D.T. Stuss, and R.T. Knight, eds. (Oxford University Press), pp. 278–291.

Duncan, J., and Miller, E.K. (2013). Adaptive neural coding in frontal and parietal cortex. In Principles of Frontal Lobe Function: Second Edition, D.T. Stuss, and R.T. Knight, eds. (Oxford University Press).

Eichenbaum, H. (2017). Barlow versus Hebb: When is it time to abandon the notion of feature detectors and adopt the cell assembly as the unit of cognition? Neurosci. Lett.

Engel, A.K., and Fries, P. (2010). Beta-band oscillations—signalling the status quo? Curr. Opin. Neurobiol. 20, 156–165.

Ferrera, V.P., Yanike, M., and Cassanello, C. (2009). Frontal eye field neurons signal changes in decision criteria. Nat. Neurosci. 12, 1458–1462.

Fitzgerald, J.K., Freedman, D.J., and Assad, J.A. (2011). Generalized associative representations in parietal cortex. Nat. Neurosci. 14, 1075–1079.

Freedman, D.J., and Assad, J.A. (2006). Experience-dependent representation of visual categories in parietal cortex. Nature 443, 85–88.

Freedman, D.J., and Assad, J.A. (2016). Neuronal mechanisms of visual categorization: an abstract view on decision making. Annu. Rev. Neurosci. 39, 129–147.

Freedman, D.J., Riesenhuber, M., Poggio, T., and Miller, E.K. (2001). Categorical representation of visual stimuli in the primate prefrontal cortex. Science 291, 312.

Freedman, D.J., Riesenhuber, M., Poggio, T., and Miller, E.K. (2003). A comparison of primate prefrontal and inferior temporal cortices during visual categorization. J. Neurosci. 23, 5235–5246.

Fries, P. (2005). A mechanism for cognitive dynamics: neuronal communication through neuronal coherence. Trends Cogn. Sci. 9, 474–480.

Fusi, S., Miller, E.K., and Rigotti, M. (2016). Why neurons mix: high dimensionality for higher cognition. Curr. Opin. Neurobiol. 37, 66–74.

Goodwin, S.J., Blackman, R.K., Sakellaridi, S., and Chafee, M.V. (2012). Executive control over cognition: stronger and earlier rule-based modulation of spatial category signals in prefrontal cortex relative to parietal cortex. J. Neurosci. 32, 3499–3515.

Hampson, R.E., Pons, T.P., Stanford, T.R., and Deadwyler, S.A. (2004). Categorization in the monkey hippocampus: a possible mechanism for encoding information into memory. Proc. Natl. Acad. Sci. U. S. A. 101, 3184–3189.

Helfrich, R.F., and Knight, R.T. (2016). Oscillatory dynamics of prefrontal cognitive control. Trends Cogn. Sci. 20, 916–930.

van Kerkoerle, T., Self, M.W., Dagnino, B., Gariel-Mathis, M.-A., Poort, J., van der Togt, C., and Roelfsema, P.R. (2014). Alpha and gamma oscillations characterize feedback and feedforward processing in monkey visual cortex. Proc. Natl. Acad. Sci. 111, 14332–14341.

Kopell, N., Whittington, M.A., and Kramer, M.A. (2011). Neuronal assembly dynamics in the beta1 frequency range permits short-term memory. Proc. Natl. Acad. Sci. 108, 3779–3784.

Llinás, R.R., Ribary, U., Jeanmonod, D., Kronberg, E., and Mitra, P.P. (1999). Thalamocortical dysrhythmia: a neurological and neuropsychiatric syndrome characterized by magnetoencephalography. Proc. Natl. Acad. Sci. 96, 15222–15227.

Lundqvist, M., Rose, J., Herman, P., Brincat, S.L., Buschman, T.J., and Miller, E.K. (2016). Gamma and Beta Bursts Underlie Working Memory. Neuron 90, 152–164.

Miller, E.K. (2000). The prefrontal cortex and cognitive control. Nat. Rev. Neurosci. 1, 59–66.

Miller, E.K., and Cohen, J.D. (2001). An integrative theory of prefrontal cortex function. Annu. Rev. Neurosci. 24, 167–202.

Miller, E.K., Freedman, D.J., and Wallis, J.D. (2002). The prefrontal cortex: categories, concepts and cognition. Philos. Trans. R. Soc. Lond. B. Biol. Sci. 357, 1123–1136.

Mitra, P.P., and Pesaran, B. (1999). Analysis of dynamic brain imaging data. Biophys. J. 76, 691–708.

Moazami-Goudarzi, M., Sarnthein, J., Michels, L., Moukhtieva, R., and Jeanmonod, D. (2008). Enhanced frontal low and high frequency power and synchronization in the resting EEG of parkinsonian patients. Neuroimage 41, 985–997.

Olejnik, S., and Algina, J. (2003). Generalized eta and omega squared statistics: measures of effect size for some common research designs. Psychol. Methods 8, 434–447.

Oostenveld, R., Fries, P., Maris, E., and Schoffelen, J.-M. (2011). FieldTrip: open source software for advanced analysis of MEG, EEG, and invasive electrophysiological data. Comput. Intell. Neurosci. 2011, 1.

Rigotti, M., Barak, O., Warden, M.R., Wang, X.-J., Daw, N.D., Miller, E.K., and Fusi, S. (2013). The importance of mixed selectivity in complex cognitive tasks. Nature 497, 585.

Roy, J.E., Riesenhuber, M., Poggio, T., and Miller, E.K. (2010). Prefrontal Cortex Activity during Flexible Categorization. J. Neurosci. 30, 8519.

Seger, C.A., and Miller, E.K. (2010). Category learning in the brain. Annu. Rev. Neurosci. 33, 203–219.

Spitzer, B., and Haegens, S. (2017). Beyond the Status Quo: A Role for Beta Oscillations in Endogenous Content (Re-) Activation. Eneuro.

Stanley, D.A., Roy, J.E., Aoi, M.C., Kopell, N.J., and Miller, E.K. (2016). Low-Beta oscillations turn up the gain during category judgments. Cereb. Cortex 1–15.

Uhlhaas, P.J., and Singer, W. (2006). Neural Synchrony in Brain Disorders: Relevance for Cognitive Dysfunctions and Pathophysiology. Neuron 52, 155–168.

Voytek, B., Kayser, A.S., Badre, D., Fegen, D., Chang, E.F., Crone, N.E., Parvizi, J., Knight, R.T., and D’esposito, M. (2015). Oscillatory dynamics coordinating human frontal networks in support of goal maintenance. Nat. Neurosci. 18, 1318–1324.

Womelsdorf, T., and Everling, S. (2015). Long-range attention networks: circuit motifs underlying endogenously controlled stimulus selection. Trends Neurosci. 38, 682–700.

Womelsdorf, T., and Fries, P. (2007). The role of neuronal synchronization in selective attention. Curr. Opin. Neurobiol. 17, 154–160.

Wong, Y.T., Fabiszak, M.M., Novikov, Y., Daw, N.D., and Pesaran, B. (2016). Coherent neuronal ensembles are rapidly recruited when making a look-reach decision. Nat.Neurosci. 19, 327.

Yuste, R. (2015). From the neuron doctrine to neural networks. Nat. Rev. Neurosci. 16, 487.

